# Reducing flight time during running decreases tibial-fibular strains: a finite element analysis

**DOI:** 10.1101/2024.10.17.618929

**Authors:** Arash Khassetarash, Benno. M. Nigg, W. Brent Edwards

## Abstract

**Purpose:** Reducing strains within the tibia and fibula during running may reduce the risk of stress fractures. We examined the effect of reduced flight time during running (i.e., grounded running) on finite-element predicted bone strains within the tibia-fibula complex.

**Methods:** Nine physically active males ran on an instrumented treadmill at 2.2 m/s using a preferred and reduced flight time technique in a randomized order. Three-dimensional force and motion capture data were recorded during running and a computed tomography image was subsequently acquired for the participant’s left leg. An inverse-dynamics-based musculoskeletal modeling workflow was used to calculate bone-on-bone contact and muscle forces during the stance phase of running. These forces served as inputs to a participant-specific finite-element model to estimate peak bone strains and strained volume (i.e., the volume of bone experiencing strains above a specific threshold) within the tibia-fibula complex.

**Results:** Guided attempts to reduce flight time was successful with an 18 ms (95% CI: 12 ms, 25 ms; p<0.001) reduction in flight time. Reducing flight time was associated with significant reductions in peak tibial/fibular strains (17% lower; 95% CI: −7.1%, −25.0%; p=0.002) and strained volume (35% lower; 95% CI: −13.57%, −50.87%; p=0.007).

**Conclusion:** We conclude that guided attempts to reduce flight time significantly reduces strains in the tibia and fibula during treadmill running at a fixed speed. These results suggest that grounded running may be a viable technique to reduce musculoskeletal loading and stress fracture risk, particularly in slow runners and those runners coming back from injury.

## Introduction

Running is one of the most accessible and popular forms of physical activity, annually adopted by over 600 million people worldwide (Wonder, n.d.). Unfortunately, running-related injuries are common, and these injuries act as barriers to achieving the many physical and mental health benefits of running. Stress fractures are particularly common amongst runners, with an incident rate of 8-18 injuries per 1000 hours of running (Videbæk et al., 2015). Stress fractures have been associated with a mechanical fatigue phenomenon, wherein cyclic submaximal loading leads to microdamage accumulation and structural failure at loading magnitudes much lower than the bone’s monotonic failure strength (Edwards, 2018). The rate of microdamage accumulation and fatigue failure in bone is heavily dependent on the resulting strain from the applied loading magnitude (Haider et al., 2021; Loundagin et al., 2021). Hence, information regarding bone strain in running is of paramount importance to evaluate the efficacy of training interventions to reduce the risk of stress fractures.

Stress fractures in runners frequently occur at the tibia (Fredericson et al., 2006; Rizzone et al., 2017), and seminal work quantified tibial bone strains *in vivo* through direct strain gauge measurement. These studies investigated differences in tibial strains in response to changes in running speed (Milgrom, 2000), grade (Burr et al., 1996), surface (Milgrom et al., 2003), and fatigue (Milgrom et al., 2007). *In vivo* strain gauge measurements are invasive and typically limited to a single anatomical location, which do not characterize the complex strain distribution experienced throughout the bone (Yang et al., 2011). For this reason, more recent studies have estimated tibial strains in different running conditions by means of musculoskeletal modeling and finite element (FE) analysis (Baggaley et al., 2024; Bruce et al., 2022; Burnet, 2017; Haider et al., 2020). In addition to a noninvasive estimation of peak strain, this approach has the added benefit of approximating the entire strain distribution within the bone, and the volume of bone experiencing high strain (i.e., above a presumed yield) is strongly associated with the number of cycles to failure (Haider et al., 2021).

Several modeling studies have investigated potential interventions to reduce tibial loads and/or strains in running. For instance, the work of Edwards et al. (2009) suggested that running with a reduced stride length may decrease bone strains and consequently, the risk of tibial stress fracture (Edwards et al., 2009); these findings were later verified through a prospective study in colligate runners (Kliethermes et al., 2021). Alternative interventions that may reduce tibial loads and thus the risk of stress fracture include running with a wider step width (Meardon and Derrick, 2014) and the use of advanced footwear technology (Werkhausen et al., 2024). Recently, Bonnaerens et al., (2022) investigated an intervention that eliminated the flight phase of running (i.e., “grounded running”; (Gatesy & Biewener, 1991)), which suggested reduced peak muscle and joint contact forces at the lower extremity with this technique. This technique may be beneficial for slower runners or those coming back from running injuries. However, given the complex relationship between applied load and bone strain, we were interested to further characterize the influence of grounded running on the mechanical environment within the tibia.

The purpose of this study was to investigate the effects of grounded running on bone strains in the tibia-fibula complex using a combination of experimentation, musculoskeletal modelling, and FE analysis. To this end, an inverse-dynamics based musculoskeletal model was used to estimate lower-extremity muscle forces while participants ran with a preferred or grounded form of running. Participant-specific FE models based on computed tomography (CT) scans were then used to compute tibial-fibular strains in each condition. Based on previous research, suggesting reduced lower-extremity joint contact forces in grounded running (Bonnaerens et al., 2022), we hypothesized that shortening the flight phase in running (indicative of grounded running) would reduce strains within the tibia-fibula complex when compared to preferred running. In addition, it may be beneficial to know what factors are associated with changes in bone strains in running to be used in future training interventions. For this reason, we also conducted a correlation analysis between running biomechanics, external and internal forces, and FE-predicted strains. We hypothesized that changes in muscle forces associated with changes in running technique would be correlated with tibial bone strain.

## Methods

Previous studies comparing the effect of preferred versus grounded running demonstrated an effect size of Cohen’s *d* = 2.75 for external forces (Bonnaerens et al., 2019) and Cohen’s *d* = 1.25 for internal forces (e.g., Soleus force (Bonnaerens et al., 2022)). Given uncertainty in the effect size for the primary outcome of bone strain, an *a priori* sample size of N = 8 was calculated, assuming a cautious effect size of Cohen’s *d* = 1.0 to reach a statistical power of 1-*β* = 0.8, at a significance level of *α* = 0.05 (GPower 3.1.9.7). To account for possible dropouts, a convenience sample of ten physically active males were recruited and only nine participants (age: 27.2 ± 1.98 years, height: 1.76 ± 0.09 m, body mass: 75.5 ± 11.5 kg) completed the entire protocol described below. The participants did not have any musculoskeletal injuries prior to data collection. The study protocol was approved by the local ethics committee (REB #19-1845) and all participants provided written informed consent.

Participants visited the lab on three different occasions with a maximum of a one-week between sessions. In the first visit, participants were familiarized with the grounded running technique. Participants ran on an instrumented treadmill (Bertec Corps., Columbus, OH), while the vertical component of the ground reaction force (VGRF) was projected in real time on a screen facing the runner. Grounded running was defined by the elimination of a flight phase and no double-leg-support (Gatesy and Biewener, 1991). This was accomplished by asking participants to remove the flight phase while keeping the lowest VGRF slightly above zero. An error threshold was identified on the screen by a ribbon signifying a VGRF below 50 N. Participants were given as much time as needed to learn and run comfortably with the grounded running technique. In the second visit, twenty retroreflective markers were used to capture the lower extremity’s motion during running. The participants ran on the instrumented treadmill with a preferred and grounded running technique for 5 minutes each in a randomized order; running velocity was held constant at 2.22 m/s for both conditions and the VGRF was projected on the screen in a similar manner to the first visit. Only one speed was investigated because maintaining a grounded running technique at faster speeds was not feasible for all participants. Three-dimensional external forces and moments from the instrumented treadmill were recorded at 2000 Hz and motion capture data were recorded at 200 Hz (Vicon motion Analysis Ltd., Oxford, UK) for 30 seconds at a 2.5-minute mark of each condition. In the third visit, a computed tomography (CT) scan was obtained using a GE Revolution GSI (GE Healthcare, Chicago, IL) with acquisition settings of 120 kVp and 180 mA. The CT scans captured the entire participants’ left leg and were reconstructed with an in-plane resolution of 0.486 mm by 0.486 mm and a slice thickness of 0.625 mm. A hydroxyapatite calibration phantom (QRM GmbH, Mohrendorf, Germany) was placed in the field of view of each scan to transform CT attenuation values in Hounsfield units to bone mineral equivalent density (*ρ*_HU_).

We used a previously established inverse-dynamics-based musculoskeletal modeling pipeline to estimate muscle and joint contact forces in the left leg during the running protocol (Baggaley et al., 2021; Edwards et al., 2009). The musculoskeletal model included 44 lower extremity muscle-tendon units based on Arnold et al. (Arnold et al., 2010). For every 1% of the gait cycle, muscle forces were estimated by minimizing the following cost function (Crowninshield and Brand, 1981):

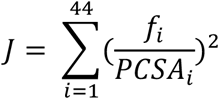

where *f*_*i*_ is the estimated force for the *i*th muscle and *PCSA*_*i*_ is the physiological cross-section of the *i*th muscle. The muscle forces were bound between zero and the maximum dynamic forces, adjusted for length and velocity. The moments produced by the muscles about the lower extremity joints were constrained to match the joint moments predicted by the inverse dynamics analysis. Specifically, the flexion-extension, and abduction-adduction moments at the hip, and the flexion-extension moment at the knee and ankle were used as constraints. The optimization was implemented using the interior-point algorithm in the MATLAB Toolbox (i.e., fmincon function). Bone-on-bone joint contact forces were calculated as the algebraic sum of muscle forces from the musculoskeletal model and intersegmental reaction forces from inverse dynamics.

FE models of the tibia-fibula complex followed the procedures previously published by our group (Baggaley et al., 2024; Khassetarash et al., 2023), which predicted tibial bending and torsion similar to those from experimental bone pin studies (Yang et al., 2014) and strains from experimental measurements (Burr et al., 1996). Briefly, the CT images were manually segmented and converted into 3D models of the tibia and fibula using the Mimics Innovation Suit (v21, Materialize, Leuven, Belgium). The 3D tibia and fibula models were then meshed using NetGen/NGSolve (https://ngsolve.org). Bones were meshed with quadratic tetrahedral elements with an average of ∼300,000 elements for the tibia and ∼50,000 elements for the fibula. A convergence study from our previous work illustrated that changing the number of elements from 130,000 to 270,000 changed peak tibial strains by 4% (Bruce et al., 2022), suggesting the FE mesh was adequately refined. Bone was modeled as an inhomogeneous orthotropic material with bone apparent density (*ρ*_*app*_) being used to assign element moduli according to:

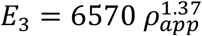

where *E*_3_ is the elastic modulus in the longitudinal direction and *ρ*_*app*_ = *ρ*_HU_ /0.626 g/cm^3^ (Dalstra et al., 1993). The elastic and shear moduli in the other directions along with the Poisson’s ratios were determined according to Rho et al., (Rho et al., 1995) as reported in our previous work (Haider et al., 2020)

Boundary conditions were defined similar to Khassetarash et al., (2023), where the tibial plateau was fixed in all degrees of freedom and a point on the medial proximal tibia was fixed in the anterior-posterior direction; the ankle joint was fixed in the anterior-posterior and medio-lateral directions. The tibia and fibula were connected by binding adjacent elements of the bones at the proximal and distal side. The anterior and the posterior proximal tibia-fibula ligaments were modelled as spring elements with stiffness according to Marchetti et al. (2017). The ankle joint contact force along with a residual moment (Haider et al., 2020) were applied to the ankle joint centre. Forces from sixteen muscles along with the patellar tendon were applied as point forces at their insertions on the tibia or fibula. The quasi-static finite element simulation was performed in Abaqus (version 2021, Dassault System, RI, USA) using muscle forces from the instant of peak ankle joint contact force. To further verify model predicted strains, similar to Baggaley et al., (2024), we created a virtual strain gauge between the midshaft and 2 cm distal on the medial tibial surface. We then compared the strains at the site of the virtual strain gauge to the experimental studies (Burr et al., 1996).

We focused our analysis on elements in the tibial and fibular diaphysis (Bruce et al., 2022). For these elements, we calculated the 90^th^ percentile pressure modified von Misses strain (de Vree et al., 1995) as a measure of peak strain and strained volume (defined as the volume of elements experiencing a pressure modified von Misses strain above 3000 με), as the latter was shown to correlate well with the fatigue life of whole rabbit tibiae (Haider et al., 2021). To quantify changes in biomechanics associated with running technique, secondary dependent variables of interest included spatiotemporal parameters (i.e., flight time, contact time, step frequency, duty factor) and the VGRF.

A Shapiro-Wilk test of normality was performed on all variables of interest. The results were reported in mean ± standard deviation unless otherwise stated. Running technique (i.e., preferred or grounded) was treated as an independent variable while the FE results (i.e., 90^th^ percentile strain and strained volume) as well as spatiotemporal and VGRF were treated as dependent variables. Paired t-tests were conducted for dependent variables to compare preferred and grounded running conditions. An exploratory correlation analysis was also performed between FE predicted strains, spatiotemporal variables, VGRF, and select muscle or tendon forces (i.e., soleus, patellar tendon). These two muscle and tendon forces were chosen because of their large magnitudes and significant contributions to the bone strain distribution. A Benjamini-Hochberg (Benjamini and Hochberg, 1995) false discovery rate correction was applied to the significance values from the correlation analysis; statistical significance was set at 0.05 after the p-values were corrected.

## Results

Attempts to decrease flight time through grounded running were successful. Indeed, flight time was reduced from 31 ± 11 ms during preferred running to 13 ± 4 ms during grounded running (*p* < 0.001) (**Figure 1**). No changes in contact time (345 ± 25 ms in preferred condition vs. 354 ± 21 ms in grounded running; P = 0.21) or step frequency (160 ± 8.8 steps per minute in preferred condition vs. 164 ± 8.9 steps per minute in grounded running; *p* = 0.24) associated with the intervention were observed. Duty factor increased from 0.46 ± 0.16 during preferred running to 0.48 ± 0.06 during grounded running (*p* < 0.001). The VGRF also reduced significantly during grounded running (1605 ± 211 N vs. 1351 ± 240 N; *p* = 0.006).

**Figure 1.**
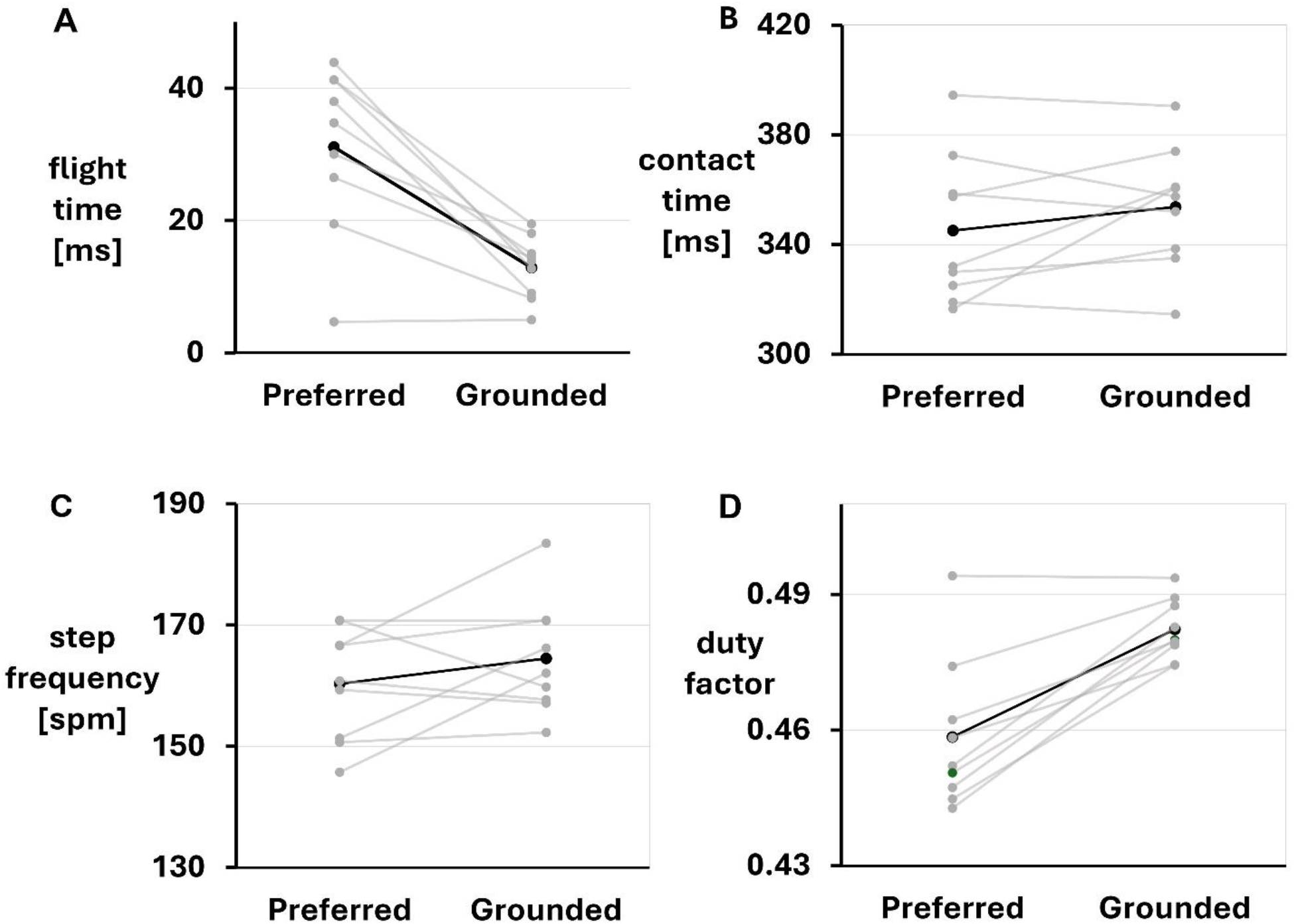
Spatiotemporal variables during preferred and grounded running including: A) flight time B) contact time, C) step frequency, and D) duty factor. In both graphs, the thick black line represents the mean of all participants, and the gray transparent lines represent individual participants.

The distribution of pressure-modified von Misses strain for a representative participant is shown in **Figure 2**. In general, the strains on the anterior and posterior aspects of the tibial and fibular midshaft were lower during grounded running when compared to preferred running. **Figure 3** depicts the strained volume for the same participant. In both conditions, most of the volume of bone experiencing over 3000 με (i.e., elements contributing to the strained volume calculation) were on the posterior aspect of the tibia and fibula. Quantitatively, peak strain was significantly lower during grounded running compared to preferred running (**Figure 4A**, 4193 ± 737 με vs. 3498 ± 738 με; *p* = 0.002). Strained volume was also significantly reduced in the grounded running condition when compared to preferred running (**Figure 4B**, 9680 ± 3159 mm^3^ vs. 6305 ± 2905 mm^3^; *p* = 0.007).

**Figure 2.**
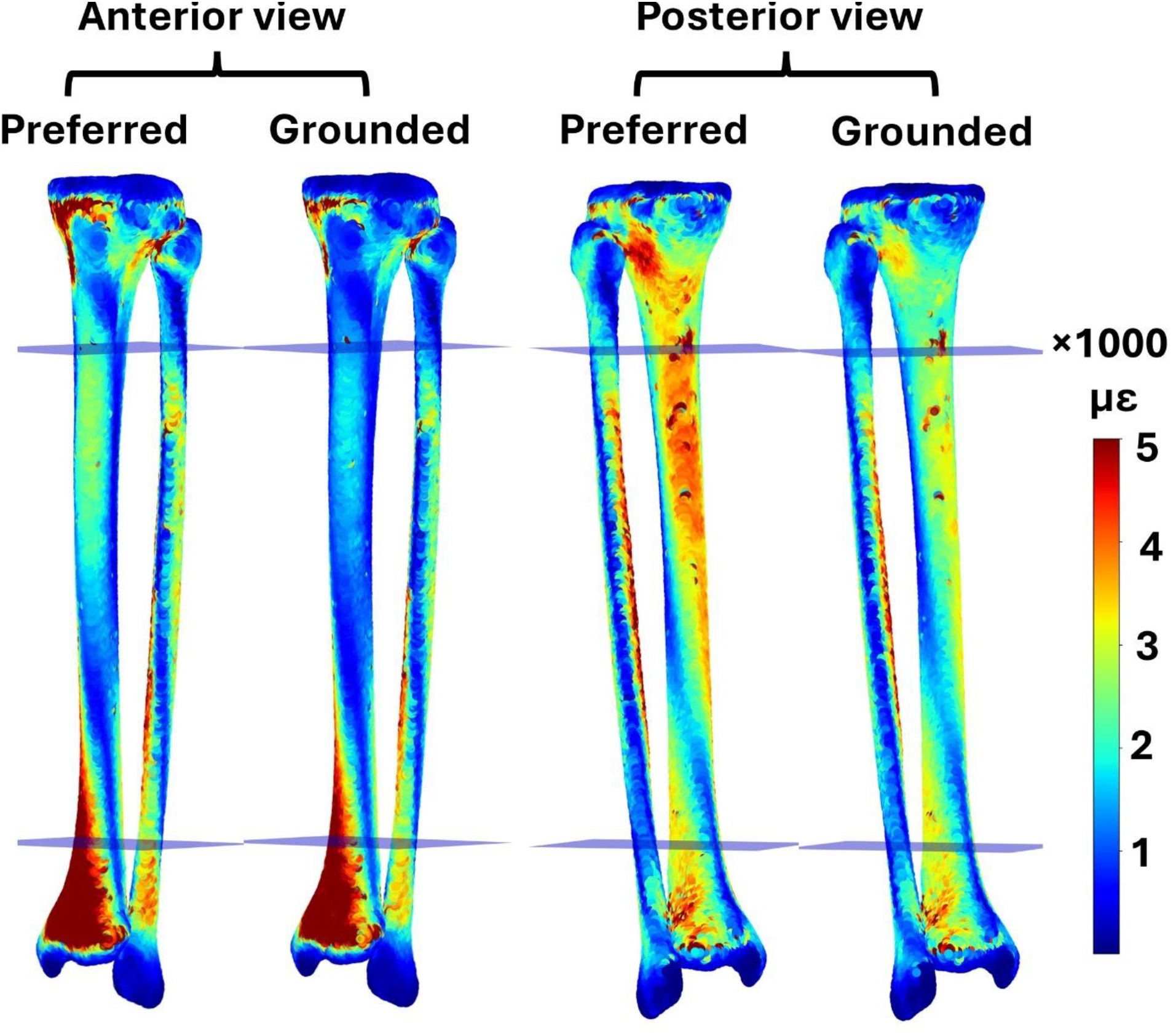
Pressure-modified von Misses strain distribution in the tibia-fibula complex during preferred and grounded running for a representative participant. The elements between the proximal and distal planes shown in the figure were considered for peak strain calculation.

**Figure 3.**
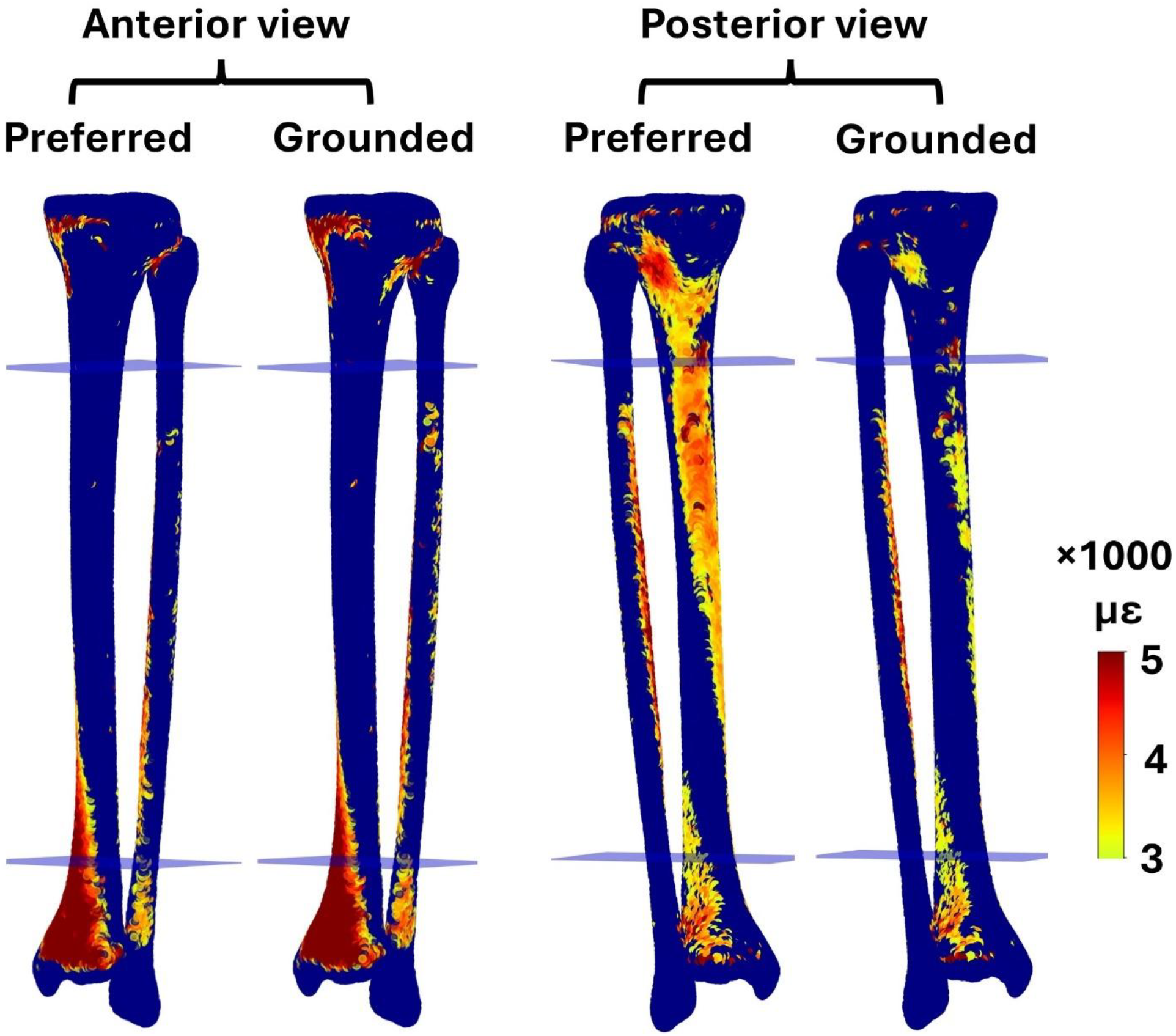
Strained volume (i.e., volume of elements experiencing strains above 3000 με) during preferred and grounded running for a representative participant. The elements between the proximal and distal planes shown in the figure were considered for strained volume calculation.

**Figure 4.**
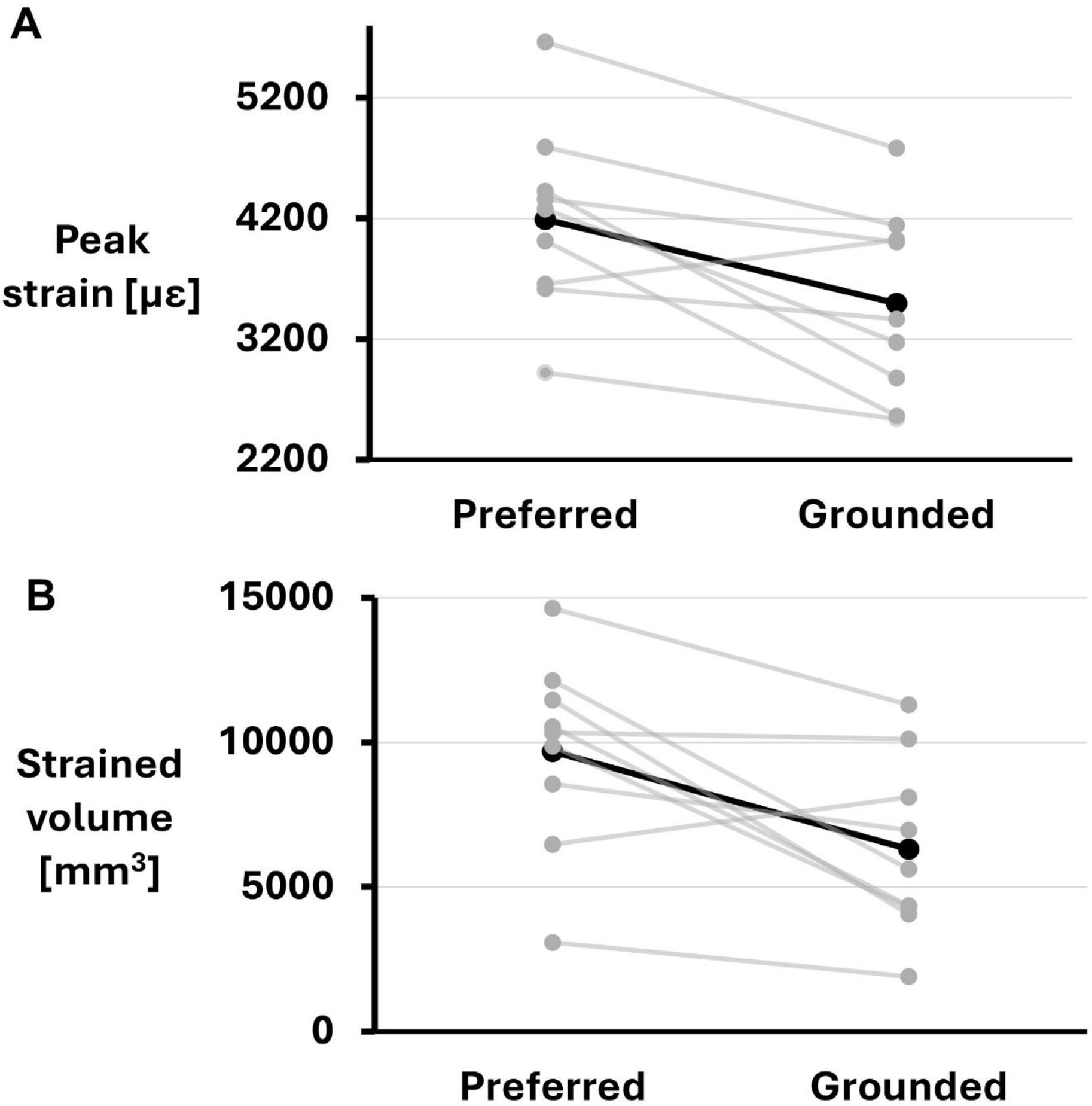
Finite element predicted A) peak strain (i.e., 90th percentile strain within the midshaft of the tibia-fibula complex) and B) strained volume (i.e., volume of bone experiencing strains above 3000 με within the midshaft of tibia-fibula complex). In both graphs, the thick black line represents the mean of all participants, and the gray transparent lines represent individual participants.

Correlation analyses (**Table 1**) revealed that between 70% to 90% of the change in peak strain could be explained by peak patellar tendon force (*p* = 0.016) and peak soleus force (*p* < 0.001), while strained volume was not well-explained by any of the variables explored (all *p ≥* 0.19).

**Table 1.** Correlation analysis between the finite element outputs and select musculoskeletal loading and running biomechanics variables.

|  | Variable | R-squared | Corrected<br>*p*-value |
| --- | --- | --- | --- |
| **Δ peak strain** | Δ VGRF | 0.160 | 0.328 |
|  | Δ flight time | 0.453 | 0.468 |
|  | Δ duty factor | 0.712 | 0.097 |
|  | Δ Soleus force | 0.912 | **<0.001** |
|  | Δ Patellar tendon force | 0.702 | **0.023** |
| **Δ strained volume** | Δ VGRF | 0.233 | 0.285 |
|  | Δ flight time | 0.317 | 0.192 |
|  | Δ duty factor | 0.388 | 0.336 |
|  | Δ Soleus force | 0.517 | 0.328 |
|  | Δ Patellar tendon force | 0.5641 | 0.192 |

The strains predicted at the virtual strain gauge site were within the range of the experimental strain gauge study. During preferred running condition, the FE model predicted maximum principal strains of 853 ± 313 με, minimum principal strains of −955 ±-454 με, and maximum shear strains of 1809 ± 522 με. These values are comparable to Burr et al. (1996) (Burr et al., 1996) who reported maximum principal strains of 625 ± 15 με, minimum principal strains of −879 ± −73 με, and maximum shear strains of 1444 ± 141 με during jogging at 2.82 m/s.

## Discussion

Mechanical fatigue is believed to play an important role in the development of stress fracture (Edwards, 2018), and the fatigue behaviour of bone is strongly dependent on the resulting strain from the applied cyclic load (Carter et al., 1981). In this study, we investigated the effects of grounded running (i.e., reducing flight time) on FE-predicted tibial/fibular strains. We observed a significant reduction in both the peak strain and strained volume in grounded running when compared to a preferred running technique. The reductions in peak strain were explained, in part, by corresponding reductions in muscle forces associated with grounded running. These findings suggest that grounded running may represent a practical gait strategy for lowering the risk of stress fractures.

The reductions in peak strain and strained volume associated with grounded running would be expected to reduce the risk of stress fracture considerably. On average, grounded running reduced the magnitude of peak strain by 695 ± 581 με (95% CI: 316 με, 1076 με) when compared to preferred running. Using a stress-life approach, we can estimate how much longer a participant could hypothetically run by adopting a grounded running technique before bone failure. The relationship between bone stress, or strain, and the number of cycles to failure is well explained by an inverse power-law with an exponent ranging from −5 to −15 (Martin, 1992). Assuming an exponent of −7, used previously for estimations of bone damage (Carter and Caler, 1985; Matijevich et al., 2019) and stress fracture risk (Edwards et al., 2009), the ∼17% reduction in peak strain associated with grounded running would correspond to a ∼256% increase in the number of cycles to failure. In other words, a grounded running technique may allow an individual to run more than three times farther at the same speed before experiencing a stress fracture. Note that this simplified estimation does not consider the large scatter observed in *ex vivo* cyclic tests of bone (Taylor, 1999), or the bone adaptation and bone remodeling response to microdamage and mechanical loading (Burr et al., 1985).

Prior research investigating grounded running demonstrated a reduction in external forces (Bonnaerens et al., 2019) as well as muscle and joint contact forces (Bonnaerens et al., 2022) when compared to a preferred technique. The current study contributes to this literature by demonstrating these reductions in applied load were also associated with concurrent reductions in bone strain. Our findings underscore the limitations of relying solely on external loading parameters for predicting bone strain. Indeed, the correlation analysis between FE outputs and VGRF revealed weak associations (R^2^ = 0.16-0.23), indicating that external forces alone are insufficient predictors of peak strain or strained volume. On the other hand, select muscle-tendon forces explained a substantial portion of the variance in peak strain (R^2^ = 0.70-0.91), suggesting their utility as a surrogate measure of bone strain and presumably stress fracture risk. These findings highlight the crucial role of muscle forces in modulating the stress-strain environment within bone, beyond what is captured by external loading variables.

While this study provides insights into the bone strain distribution in grounded running, it is important to acknowledge certain limitations inherent to musculoskeletal and FE modeling. For example, the estimation of muscle forces relied on an optimization approach with a cost function. While this cost function has been implicitly tested (Zargham et al., 2019), it lacks explicit validation. Moreover, the muscle parameters used in the model, including origins, insertions, moment arms, and theoretical maximum forces, were based on estimations rather than direct measurements from our participants. These estimations may affect the magnitude of strain but are highly unlikely to affect our interpretation given the repeated measure design of this study. With regard to the FE model, while the boundary conditions were grounded in physiological principles (Haider et al., 2020), they may not perfectly replicate the actual joint mechanical behavior *in vivo*. Nevertheless, FE predicted bone strains at a virtual strain gauge location were similar to those measured experimentally by Burr et al., (Burr et al., 1996). The peak strains in our FE models were considerably higher (ranging from 3000 με to 5000 με) than those at the virtual strain gauge location, which illustrates the shortcomings of using a single strain gauge measurement to characterize the complex strain distribution experienced throughout the bone. Despite uncertainties related to the musculoskeletal and FE modeling approach used herein, our current and prior research (Baggaley et al., 2024; Khassetarash et al., 2023) has demonstrated promising results in predicting bone bending and strains that align with experimental findings from bone pin (Yang et al., 2014) and strain gauge (Burr et al., 1996; Milgrom et al., 2020) studies.

It is also crucial to remember that this investigation focused on an acute intervention, and the long-term implications of prolonged grounded running on the musculoskeletal system remain unknown. Future studies should consider the potential for adaptations and changes in bone strains with chronic exposure to grounded running. Grounded running was accomplished with minimal instruction, but this was facilitated through real-time visual feedback. If grounded running program were to be implemented in practice several laboratory-based gait training sessions would likely be required (e.g., (Gaudette et al., 2022)). In this regard, there are several questions left to be answered related to the length and frequency of gait training sessions for optimal adoption and recall. Although we would not recommend grounded running as a universal strategy, in part because of the slow running speed required, it could represent a viable load-management technique for those runners with a history of overuse injury, particularly in situations where running speed can be sacrificed.

## Conclusion

This study suggested that reducing the flight phase of running by transitioning to a grounded running technique is an effective intervention for decreasing strains within the tibia-fibula complex in the majority of runners. Although confirmation through prospective studies is still needed, reducing the flight phase appears to be a promising strategy to reduce bone strain and stress fracture risk at slower running velocities.

## Acknowledgment

We acknowledge the support of the Natural Sciences and Engineering Research Council of Canada (<u>RGPIN 02404-2021</u>). We would like to thank Art Kuo, Jeremy Wong, Koen Lemire for their help in study design and implementation, and Mark Pineda for the help with data collection.

## Conflict of interest

No conflict of interest to declare.

## Notes

### Competing Interest Statement

The authors have declared no competing interest.

### Summary of Updates

We have added a section to calculate the planar strains at a virtual strain gauge location around tibial midshaft and compared the results to experimental work by Burr et al., 1996. This comparison further strengthens and validates the results of our finite element analysis.

